# Evaluating the position of the uncal apex as a predictor of episodic memory across the adult lifespan

**DOI:** 10.1101/2024.03.18.585629

**Authors:** Philip Bahrd, Micael Andersson, Grégoria Kalpouzos, Alireza Salami, Kristin Nordin

**Author notes:** Current address: German Center for Neurodegenerative Diseases, Magdeburg, Germany (Philip Bahrd). Corresponding author: Kristin Nordin Aging Research Center Karolinska Institutet Tomtebodavägen 18A 171 65 Solna, Sweden.

## Abstract

Structural decline of the hippocampus follows heterogeneous spatial patterns, and is an important determinant of episodic memory dysfunction in aging. However, evidence indicate that the anatomical landmark uncal apex, used to demarcate anterior and posterior hippocampal subregions, changes position as the hippocampus atrophies. This risks misclassifying gray matter into the incorrect subregion when using standard demarcation methods, potentially contributing to over– and underestimation of age-related anterior and posterior hippocampal volume loss, respectively. Yet, it remains unexplored whether inter-individual differences in uncal apex position itself predict episodic memory performance. Here, we manually identified the uncal apex in anatomical MRI data from a healthy adult-lifespan sample (n=180; 20-79 years), analyzed age differences in its position, and examined associations with word recall performance. Increasing age was linked to a more anteriorly located uncal apex (retracting ∼0.04 mm/year), and this anterior shift was linked to lower memory performance. Using the manually identified uncal apex for subregion segmentation in native space revealed a lower proportion of anterior hippocampal volume compared to segmentation in MNI space (y = –21), with effects of age on volumes differing between methods. Memory performance was not predicted by hippocampal subregion volumes, but the ratio of posterior to anterior volume was significantly linked to memory when accounting for tissue misclassification due to uncal apex variation. These results indicate that uncal apex position may provide an estimate of hippocampal integrity sensitive to inter-individual differences in memory, independent of limitations associated with different segmentation methods.

## 1. Introduction

Age-related decline in episodic memory has several biological sources, among them changes in the structural integrity of the hippocampus (Fjell et al., 2014; Gorbach et al., 2017). Consistent with the anatomical and functional heterogeneity characterizing the hippocampus longitudinal axis (Poppenk et al., 2013; Strange et al., 2014), previous studies have demonstrated value in assessing gray matter atrophy and associations with memory separately for anterior and posterior hippocampal subregions (Chauveau et al., 2021; Langnes et al., 2020; Malykhin et al., 2017; Nordin et al., 2018; Snytte et al., 2022). However, a previous study observed that the position of the anatomical landmark most commonly used for segmenting the hippocampus into anterior and posterior subregions – the uncal apex – varies as a function of age, increasing the risk of misclassifying anterior hippocampal tissue as part of the posterior hippocampus in an age-dependent manner (Poppenk, 2020).

Using cross-sectional data from several large-scale samples, Poppenk (2020) demonstrated an anterior displacement of the uncal apex by approximately 0.03 mm/year across the adult lifespan, and by 0.05-0.06 mm/year in older age (>60 years). Whereas this observation has limited implications for segmentation of the anterior and posterior hippocampus in healthy young adults, it suggests that an uncal apex landmark-based approach may be inappropriate for demarcating anterior and posterior subregions in samples where anatomical hippocampal alterations occur. The current understanding posits that, as the uncal apex retracts anteriorly, it exposes the sub-uncal area – which in older and some clinical samples would risk being misclassified as posterior tissue (whereas still accurately classified as anterior tissue in healthy young adults). In line with this, Poppenk (2020) reported overestimated anterior, while underestimated posterior, volume loss when volumes were based on uncal apex landmark-segmentation as compared to when based on segmentation in normalized space at the Montreal Neurological Institute (MNI) coordinate of y = –21. Critically, age-related variation in the position of the uncal apex would risk misclassification of any hippocampal feature estimated using anterior and posterior hippocampal masks (e.g., gray matter, functional activation, connectivity, molecular markers).

While previous research has established significant heterogeneity in associations with episodic memory between anterior and posterior hippocampal subregions, findings remain inconsistent. Whereas a recent large-scale longitudinal lifespan study indicated a primary link between anterior hippocampal volume and verbal recollection (Langnes et al., 2020), cross-sectional data from the adult lifespan, in contrast, have demonstrated a main role of the posterior hippocampus in verbal recollection memory (Hoang et al., 2024; Malykhin et al., 2024), an observation consistent with an anterior-posterior division between longitudinal decline in recognition and recollection memory, respectively (Persson & Andersson, 2022). In young adults, the ratio of posterior-to-anterior volume has successfully been used to predict recollection memory (Poppenk & Moscovitch, 2011; Snytte et al., 2020), but less is known about this measure as a predictor of memory in older samples. Since this measure is directly linked to the position of the uncal apex, and therefore susceptible to any age-related differences therein, it should warrant particular consideration of inter-individual differences in uncal apex position when used in aging samples.

In replicating the methods of Poppenk (2020) to establish the impact of uncal apex position on hippocampal volumetric estimates across the adult lifespan, the specific aim of this study was to evaluate individual differences in uncal apex position as a hippocampus-based predictor of episodic memory across the adult lifespan, avoiding limitations associated with demarcating anterior and posterior subregions for volumetric estimation. We predicted that a more anteriorly located uncal apex, likely conveying reduced hippocampal structural integrity, would be linked to lower levels of memory function. Furthermore, we assessed the impact of potential tissue misclassification on associations of anterior and posterior hippocampal volume with memory by testing volumes from both landmark-based and MNI y –21 coordinate-based segmentation.

## 2. Materials and methods

This study included data from the DopamiNe, Age, connectoMe, and Cognition (DyNAMiC) study, which has been described in detail elsewhere (Johansson et al., 2023; Nordin et al., 2022). Here, we include the materials and methods directly relevant to the current study. The DyNAMiC study was approved by the Regional Ethical board of Umeå, Sweden. All participants provided written informed consent prior to testing.

### 2.1 Participants

The initial DyNAMiC sample included 180 participants (20-79 years; mean age = 49.8±17.4; 90 men and 90 women equally distributed within each decade). Recruitment was done via postal mail to a randomly selected sample from the population register of Umeå in northern Sweden. Exclusion criteria implemented during the recruitment procedure included brain pathology, impaired cognitive functioning (Mini Mental State Examination < 26), medical conditions and treatment that could affect brain functioning and cognition (e.g., dementia, diabetes, and psychiatric diagnosis), and magnetic resonance imaging (MRI) contraindications (e.g., metal implants). All participants were native Swedish speakers.

### 2.2 Episodic memory

Episodic memory was assessed using three tasks that measured word recall, number-word recall, and object-location recall (Nevalainen et al., 2015; Nordin et al., 2022). In the word recall task, participants were shown 16 Swedish concrete nouns, presented one at a time on a computer screen. Each word was displayed for 6 seconds, with an inter-stimulus interval (ISI) of 1 second. After the encoding phase, participants attempted to recall and type as many words as possible. This task was conducted over two trials, with a maximum possible score of 32. The number-word recall task required participants to memorize pairs of two-digit numbers and concrete plural nouns (e.g., 46 dogs). During the encoding phase, eight number-word pairs were displayed, each for 6 seconds, with an ISI of 1 second. In the recall phase, the nouns were presented in a shuffled order, and participants had to recall and input the corresponding two-digit number for each noun (e.g., How many dogs?). This task also consisted of two trials, with a maximum possible score of 16. The third task assessed object-location memory. Participants were shown a 6 × 6 square grid where 12 objects were sequentially placed in distinct locations. Each object was displayed in its location for 8 seconds, with an ISI of 1 second. In the recall phase, all 12 objects appeared next to the grid, and participants had to reposition them in their correct locations within the grid. If they were unsure, they were instructed to make their best guess. This task also included two trials, with a maximum possible score of 24.

A composite episodic memory measure was calculated by, for each of the three tasks, summarizing scores across the total number of trials. The three resulting sum scores were z-standardized and averaged to form one composite score of episodic memory performance (T score: mean = 50; SD = 10). Missing values were replaced by the average of the available observed scores. Our main analyses focused on composite episodic memory and, specifically, word recall performance to enhance comparability with previous research on verbal memory (Langnes et al., 2020; Persson & Andersson, 2022; Poppenk & Moscovitch, 2011). Number-word and object-location recall analyses are provided in the Supporting Information.

### 2.3 Structural MR Imaging

Brain imaging was conducted at Umeå University Hospital, Sweden. Structural MRI data were acquired with a 3T Discovery MR 750 scanner (General Electric, WI, USA), using a 32-channel head coil. Anatomical T1-weighted images were acquired with a 3D fast-spoiled gradient-echo sequence (176 sagittal slices with a thickness of 1 mm, repetition time (TR) = 8.2 ms, echo-time (TE) = 3.2 ms, flip angle = 12°, field of view (FOV) = 250 × 250 mm, in-plane resolution = 0.49 x 0.49 mm).

### 2.4 Manual identification of the uncal apex

The uncal apex, defined as the most posterior coronal slice in which the uncus is visible (Dalton et al., 2017; Malykhin et al., 2007; Poppenk, 2020), was for each participant manually identified in left and right hemispheres using anatomical T1-weighted images. To ensure reliability of manual ratings, two independent raters underwent training on how to recognize the landmark and established a common protocol. The raters independently completed an initial round of manual identification using a set of 10 randomly selected participants (i.e., 20 hippocampi). Reliability of ratings in this first round was determined as high based on an inter-class coefficient (ICC) at 0.986, which increased to 0.996 after a second round was completed. Subsequently, one of the raters (P.B.) completed the manual landmark identification in the rest of the DyNAMiC sample. Cases marked as difficult during this procedure were revisited and discussed before a final coordinate was determined.

Following the methods of the original report (Poppenk, 2020), the uncal apex coordinates identified in native space were, for each participant and hemisphere, transformed to standard MNI space using linear affine normalization (here, in Statistical Parametric Mapping, SPM12: Wellcome Trust Centre for Neuroimaging http://www.fil.ion.ucl.ac.uk/spm/). The resulting left– and right-hemisphere y-plane MNI coordinates were taken to denote *uncal apex position* (Figure 1A). Manual identification resulted in estimates of uncal apex position for a total of 173 participants (left hemisphere: M = –19.75, SD = 2.88; right hemisphere: M = –19.22, SD = 2.73), excluding 7 individuals due to problematic ratings and outlier values (1.5 x interquartile range) following normalization.

**Figure 1.**
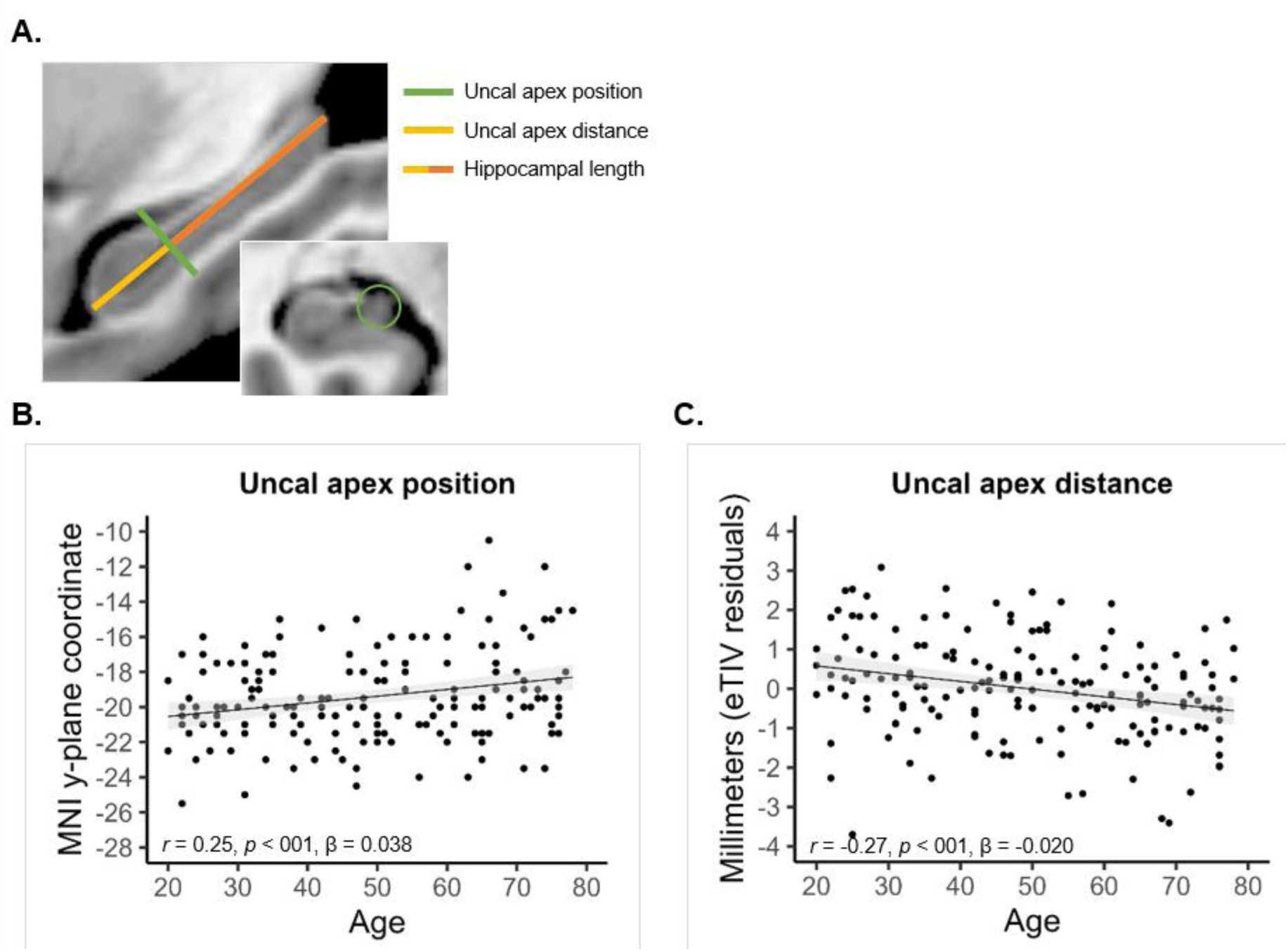
Uncal apex position– and distance estimates across the adult lifespan. A) Schematic description of the two uncal apex measures displayed on a standard MNI-normalized brain; B) Age-related differences in uncal apex position (MNI coordinates), indicating anterior displacement with increasing age; C) Age-related decrease of distance (mm, adjusted for eTIV) between the most anterior point of the hippocampus and the manually identified uncal apex.

### 2.5 Hippocampal segmentation and volumetric assessments

Individual anatomical T1-weighted images were submitted to automated segmentation in FreeSurfer version 6 (Fischl et al., 2002, 2004). Anterior and posterior hippocampal volumes were then estimated in two ways: 1) from anterior and posterior segments defined in native space by projecting the manually identified uncal apex coordinate onto an axis drawn from the most anterior to the most posterior point of the hippocampus; 2) in normalized MNI-space from anterior and posterior segments defined using the MNI y-coordinate = –21 (Poppenk et al., 2013). Volumetric values were corrected for FreeSurfer’s estimated total intracranial volume (eTIV; the sum of volumes for gray matter, white matter, and cerebrospinal fluid). Adjusted volumes were equal to the raw volume – 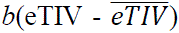, where *b* is the regression slope of volume on eTIV (Buckner et al., 2004; Jack et al., 1989). One participant was excluded based on problematic segmentation.

### 2.6 Hippocampal length and native space uncal apex distance estimates

Using the subject-specific segmentations from FreeSurfer, we additionally calculated the total length of the hippocampus and the distance in mm between the most anterior point of the hippocampus and the position of the manually identified uncal apex projected on the axis between the most anterior and most posterior points of the hippocampus. This was done to obtain an estimate of uncal apex location independent of normalization to standard space, henceforth referred to as *uncal apex distance* (Figure 1A; left hemisphere: M = 19.57 mm, SD = 1.55; right hemisphere: M = 19.07 mm, SD = 1.52). A total of 3 participants were excluded at this step due to problematic distance estimation, resulting in 169 remaining participants.

### 2.7 Statistical analyses

Statistical analyses were conducted in SPSS version 28.0 and in R version 4.1.3 (R Core Team, 2021). We considered both linear and non-linear associations between age and hippocampal measures (i.e., uncal apex position, distance, and hippocampal gray matter volume) using data-driven generalized additive models (GAM), which delineate the best fit of the data without any *a priori* assumptions of linearity. With GAM, the smoothness of the age curve is estimated as part of the model fit and allows for the effective degrees of freedom (edf) to be taken as an estimate of deviation from linearity (Zuur et al., 2009). We used simple linear regression to estimate the average annual displacement of the uncal apex and difference in volume where significant effects of age were observed. Hierarchical regression analyses were used to test uncal apex estimates and hippocampal volume as predictors of episodic memory performance over covariates (i.e., sex, age). In R, GAM was performed with the mgcv package version 1.8-40 (Wood, 2011). Statistical significance is reported using both unadjusted p-values and p-values adjusted for false discovery rate (FDR).

## 3. Results

### 3.1 Age-related anterior retraction of the uncal apex

#### 3.1.1 Uncal apex position

There was a significant correlation between uncal apex position in left and right hemispheres (*r* = 0.80, *p* < 0.001, *p*_FDR_ = 0.003), on par with previous observations in a similar lifespan sample (Poppenk, 2020: *r* = 0.72). This allowed us to combine left– and right-hemisphere y-plane MNI coordinates into a measure of mean uncal apex position (M = –19.43, SD = 2.61). GAM revealed that uncal apex position was significantly related to age across the sample (F = 11.38, edf = 1.00, *p* < 0.001, *p*_FDR_ = 0.003). Simple linear regressions estimated that the uncal apex was on average located 0.038 mm more anteriorly per year across the sample (R = 0.246, R^2^ = 0.061, β = 0.038), Figure 1B. For consistency with the previous report (Poppenk, 2020), we also estimated the effect of age across segments of individuals below and above 40 years separately. Uncal apex position did not show significant age-related variation before the age of 40 years (R = 0.167, R^2^ = 0.028, F_(1, 54)_ = 1.55, β = 0.067, *p* = 0.219, *p*_FDR_ = 0.318), but a significant average anterior displacement by 0.061 mm after the age of 40 (R = 0.250, R^2^ = 0.063, F_(1, 112)_ = 7.49, β = 0.061, *p* = 0.007, *p*_FDR_ = 0.017). Specifically, pairwise comparisons demonstrated a significantly more anterior uncal apex in older individuals (≥60 years: M = – 18.54, SD = 3.08) compared to both young (20-39 years: M = –19.95, SD = 2.25; *t* = –2.761, *p* = 0.007, *p*_FDR_ = 0.017) and middle-aged individuals (40-59 years: M = –19.80, SD = 2.21; *t* = – 2.498, *p* = 0.014, *p*_FDR_ = 0.030), who displayed equal estimates of uncal apex position (*t* = – 0.353, *p* = 0.725, *p*_FDR_ = 0.780).

#### 3.1.2 Uncal apex distance

As a native-space alternative to the MNI y-plane coordinates representing uncal apex position, we used the distance in mm from the most anterior point of the hippocampus to the manually identified uncal apex. This measure was significantly correlated between hemispheres (*r* = 0.38, *p* < 0.001, *p*_FDR_ = 0.003), and an average bilateral measure was created (M = 19.29 mm, SD = 1.30). Furthermore, estimates of uncal apex distance (controlled for eTIV to adjust for inter-individual differences in brain size) were negatively correlated with uncal apex position (*r* = – 0.35, *p* < 0.001, *p*_FDR_ = 0.003), such that a shorter distance between the hippocampus most anterior point to the uncal apex was linked to a more anterior uncal apex y-coordinate.

Uncal apex distance (estimates adjusted for eTIV) was significantly related to age across the sample (F = 12.85, edf = 1.00, *p* < 0.001, *p*_FDR_ = 0.003), such that increasing age was linked to a shorter distance between the hippocampus most anterior point and the uncal apex (Figure 1C). A linear regression estimated the annual difference in uncal apex distance to be approximately 0.020 mm (R = 0.267, R^2^ = 0.071, β = –0.020). When estimated across segments of individuals below and above 40 years separately, uncal apex distance did not show significant age-related variation before the age of 40 years (R = 0.047, R^2^ = 0.002, F_(1, 54)_ = 0.12, β = –0.011, *p* = 0.730, *p*_FDR_ = 0.780), but an average anterior displacement by 0.022 mm after the age of 40 (R = 0.207, R^2^ = 0.043, F_(1, 112)_ = 4.98, β = –0.022, *p* = 0.028, *p*_FDR_ = 0.057). Pairwise comparisons yielded a significant difference in uncal apex distance between older (M = 18.80, SD = 1.21) and young adults (M = 19.75, SD = 1.35; *t* = 3.328, *p* = 0.001, *p*_FDR_ = 0.003) and at trend-level with middle aged adults (M = 19.33, SD = 1.16; *t* = 1.966, *p* = 0.052, *p*_FDR_ = 0.091), whereas there was no difference between middle aged and young adults (*t* = –1.537, *p* = 0.127, *p*_FDR_ = 0.206).

We also examined the association between whole hippocampal length (adjusted for eTIV) and age, observing significant shortening of the hippocampus (F = 4.85, edf = 4.18, *p* < 0.001, *p*_FDR_ = 0.003). A linear model estimated the decrease in length to be on average 0.031 mm per year (R = 0.300, R^2^ = 0.090, β = –0.031).

### 3.2 Effect of uncal apex position on anterior and posterior hippocampal volumes

For comparison with previous findings (Poppenk, 2020), we estimated the effect of variation in uncal apex position on anterior and posterior hippocampal volume. Uncal apex position was negatively correlated with anterior hippocampal volume (*r* = –0.197, *p* = 0.011, *p*_FDR_ = 0.025), while positively correlated with posterior hippocampal volume (*r* = 0.169, *p* = 0.029, *p*_FDR_ = 0.057), following landmark-based segmentation in native space. On average, anterior volume decreased by 21.4 mm^3^ with every 1 mm of anterior uncal apex displacement. As such, the average difference of 1.4 mm in uncal apex position between young and older adults, indicates that misclassification due to uncal apex variation would amount to approximately 30 mm^3^ of anterior tissue (∼1% of anterior hippocampal volume).

Estimates of anterior and posterior hippocampal gray matter volume were overall greater when using the uncal apex landmark-based segmentation approach compared to coordinate-based segmentation in MNI space (anterior volume: *t* = 33.32, *p* < 0.001, *p*_FDR_ = 0. 0.003; posterior volume: *t* = 34.08, *p* < 0.001, *p*_FDR_ = 0.002; Table 1), but the proportion of anterior volume was, however, significantly lower following landmark-based (57%) compared to coordinate-based (60%) segmentation (*t* = 7.96, *p* < 0.001, *p*_FDR_ = 0. 0.003). We computed the difference in proportion of anterior hippocampal volume between segmentation methods for each individual. Linear regression analyses, controlling for age and sex, confirmed that a more anterior uncal apex was liked to a greater difference in anterior hippocampal proportion between segmentation methods, conveyed by both its position (R = 0.362, R^2^ = 0.131, β = 0.004; *p* < 0.001, *p*_FDR_ = 0. 0.003) and distance (R = 0.610, R^2^ = 0.372, β = –0.020; *p* < 0.001, *p*_FDR_ = 0. 0.003).

**Table 1.** Anterior and posterior hippocampal volumes (in mm^3^) across segmentation methods.

|  | Anterior HC |  | Posterior HC |  | % Anterior HC |
| --- | --- | --- | --- | --- | --- |
| Landmark (native space) | 2428.17 | ±274.16 | 1802.31 | ±258.09 | 57 |
| Coordinate (MNI space) | 1863.22 | ±134.28 | 1241.03 | ±106.55 | 60 |

### 3.3 Effect of segmentation method on age-related volume loss

#### 3.3.1 Age-effects on volumes from uncal apex landmark-based segmentation

Anterior hippocampal volume remained stable with increasing age (F = 0.08, edf = 1.0, *p* = 0.782, *p*_FDR_ = 0.817; R = 0.022, R^2^ = 0.000, β = –0.346), Figure 2A. In contrast, posterior hippocampal volume significantly decreased with age across the adult lifespan (F = 15.26, edf = 1.92, *p* < 0.001, *p*_FDR_ < 0.001), with an average of 6.17 mm^3^ per year (R = 0.409, R^2^ = 0.167, β = –6.171). This effect showed moderate signs of non-linearity, with an approximate onset of steeper decline at the age of 40 years (Figure 2A). To account for the degree of tissue misclassification due to uncal apex displacement, we assessed effects of age on anterior and posterior volumes controlling for the difference in proportion of anterior volume between segmentation methods. This yielded significant volume decline in both regions, now with an increased effect in the anterior hippocampus (F = 8.17, edf = 1.63, R = 0.290, R^2^ = 0.084, β = – 3.533, *p* < 0.001, *p*_FDR_ = 0.003), paralleled by a decreased effect in the posterior hippocampus (F = 12.62, edf = 1.53, R = 0.334, R^2^ = 0.111, β = –3.344, *p* < 0.001, *p*_FDR_ < 0.001).

**Figure 2.**
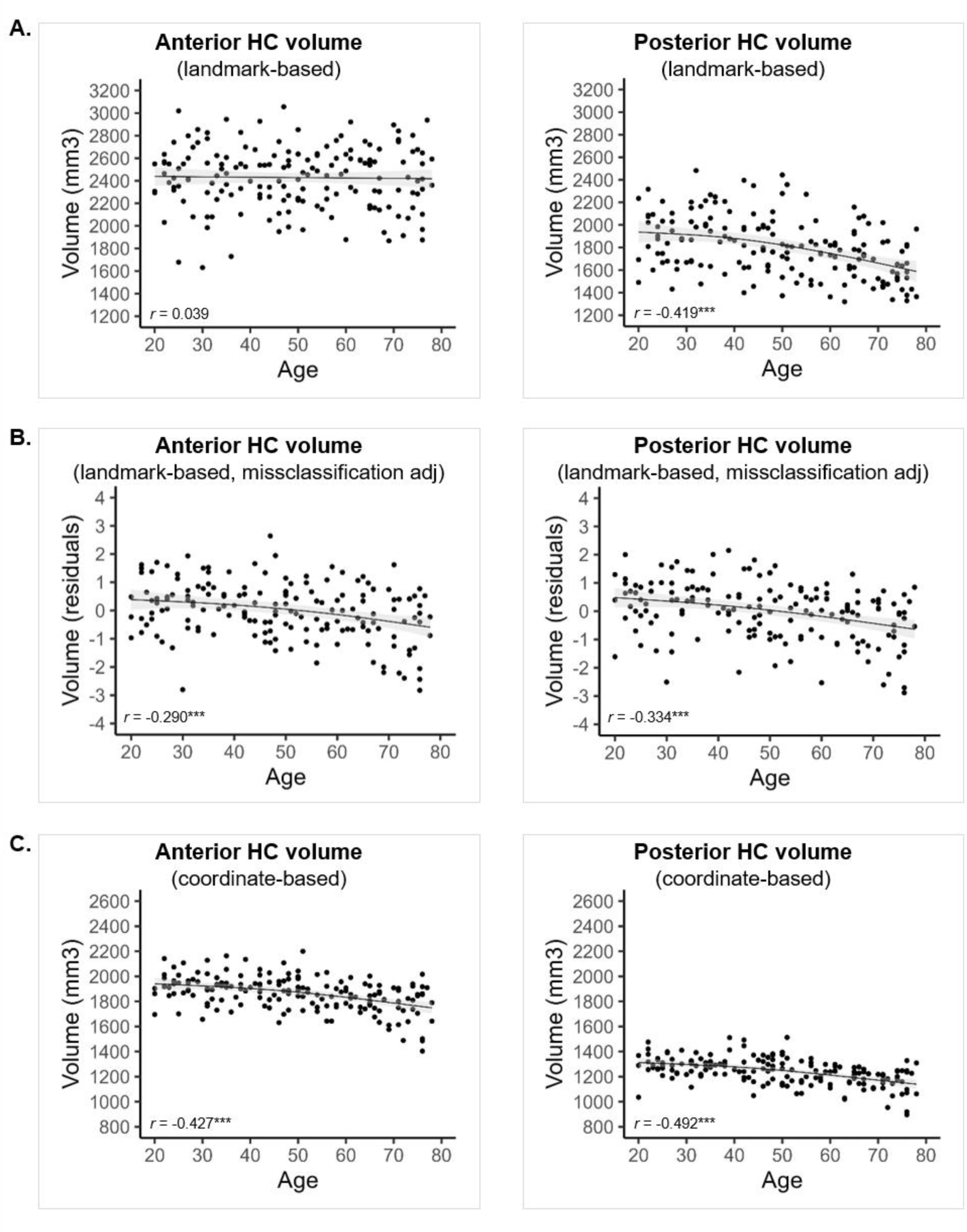
Effects of age on anterior and posterior hippocampal volume following. A) landmark-based segmentation; B) landmark-based segmentation with volumes adjusted for estimated tissue misclassification; C) coordinate-based segmentation.

#### 3.3.2 Age-effects on volumes from MNI coordinate-based segmentation

Following coordinate-based segmentation of hippocampal subregions at y = –21 in MNI-space, both the anterior and the posterior hippocampus displayed significant and non-linear negative effects of age. Consistent with the effects observed in landmark-based volumes after controlling for tissue misclassification, anterior volume decreased with an average of 3.35 mm^3^ per year (F = 17.31, edf = 1.85, *p* < 0.001, *p*_FDR_ < 0.001; R = 0.427, R^2^ = 0.182, β = –3.349) and posterior volume with an average of 3.06 mm^3^ per year (F = 23.5, edf = 1.94, *p* < 0.001, *p*_FDR_ < 0.001; R = 0.492, R^2^ = 0.242, β = –3.063). For both regions, greater decline was slightly more evident after the age of 55 years. Figure 2C.

Given the absence of the expected over-estimation of anterior volume loss by landmark-based segmentation, we conducted supplementary analyses specifically testing effects of segmentation method on anterior hippocampal volume across subgroups of older adults classified based on uncal apex position. This revealed a pattern in line with predictions, although not significant, such that older adults with the most anterior uncal apex (≥ y –17.5) showed lower anterior volume compared to the other subgroups only following landmark-based segmentation (Supporting Information Figure 1).

### 3.4 Predicting episodic memory performance

#### 3.4.1 Uncal apex position and distance

Hierarchical regression analyses were used to evaluate uncal apex position and distance as predictors of episodic memory over and above age and sex. Uncal apex position significantly predicted episodic memory measured by the composite score (ΔR^2^ = 0.016, F_(1, 166)_ = 3.98, β = –0.401, partial Pearson’s *r* = –0.15, *p* = 0.048, *p*_FDR_ = 0.087), such that a more anteriorly located uncal apex was associated with lower performance, Figure 3A. Uncal apex position similarly predicted word recall (ΔR^2^ = 0.027, F_(1, 166)_ = 7.07, β = –0.651, partial Pearson’s *r* = –0.21, *p* = 0.007, *p*_FDR_ = 0.017), Figure 3B. In contrast, uncal apex distance did not predict either composite episodic memory (ΔR^2^ = 0.006, F_(1, 166)_ = 1.57, β = 0.531, partial Pearson’s *r* = 0.10, *p* = 0.212, *p*_FDR_ = 0.318), or word recall (ΔR^2^ = 0.003, F_(1, 166)_ = 0.70, β = 0.465, partial Pearson’s *r* = 0.06, *p* = 0.422, *p*_FDR_ = 0.551).

**Figure 3.**
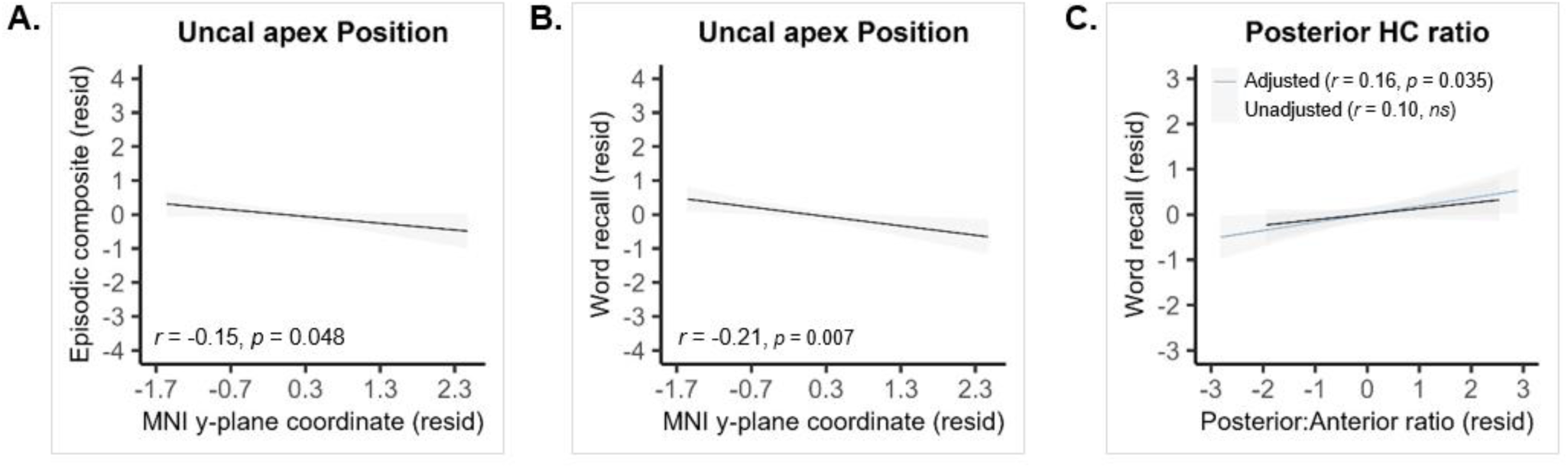
Partial regression plots (controlling for age and sex) of. A) uncal apex position as a predictor of composite episodic memory; B) uncal apex position as a predictor of word recall; and C) the ratio of landmark-based posterior to anterior volume as a predictor of word recall, with (blue) and without (black), controlling for the estimated degree of tissue misclassification.

#### 3.4.2 Anterior and posterior hippocampal volume

Overall, neither landmark-based nor coordinate-based hippocampal volumes predicted memory performance for either the composite or word recall measures. See Supporting Table 1 for all partial correlations.

#### 3.4.3. Ratio of anterior to posterior volume

The ratio of posterior to anterior volume was computed for estimates from each segmentation method, for each participant. Composite episodic memory performance was not predicted by the ratio of posterior to anterior volume, for either landmark-based (ΔR^2^ = 0.000, F_(1, 166)_ = 0.003, β = –0.20, partial Pearson’s *r* = 0.004, *p* = 0.954, *p*_FDR_ = 0.954) or coordinate-based estimates (ΔR^2^ = 0.001, F_(1, 166)_ = 0.22, β = 6.10, partial Pearson’s *r* = 0.036, *p* = 0.641, *p*_FDR_ = 0.751). Similarly, ratios from neither segmentation method predicted word recall (landmark-based: ΔR^2^ = 0.006, F_(1, 166)_ = 1.50, β = 5.10, partial Pearson’s *r* = 0.095, *p* = 0.223, *p*_FDR_ = 0.318; coordinate-based: ΔR^2^ = 0.011, F_(1, 166)_ = 2.63, β = 26.02, partial Pearson’s *r* = 0.125, *p* = 0.107, *p*_FDR_ = 0.180). Finally, we assessed the landmark-based ratio measure as a predictor of composite episodic memory and word recall when using the proportion of anterior volume difference score as a covariate. This showed that the posterior to anterior volume ratio significantly predicted word recall (ΔR^2^ = 0.018, F_(1, 164)_ = 4.52, β = 28.65, partial Pearson’s *r* = 0.164, *p* = 0.035, *p*_FDR_ = 0.066), but not episodic memory (ΔR^2^ = 0.002, F_(1, 164)_ = 0.42, β = 7.17, partial Pearson’s *r* = 0.051, *p* = 0.517, *p*_FDR_ = 0.639) when accounting for the estimated degree of tissue misclassification, Figure 3C.

## 4. Discussion

The primary aim of this study was to evaluate the position of the uncal apex as a hippocampal predictor of episodic memory across the healthy adult lifespan, and to characterize the impact of inter-individual differences in uncal apex position on anterior and posterior hippocampal volume as it relates to age and episodic memory.

The uncal apex is the anatomical landmark most commonly used to demarcate the border between the anterior and the posterior hippocampus (Dalton et al., 2017; Duvernoy, 2013; Malykhin et al., 2007; Olsen et al., 2013), but the discovery that it shifts anteriorly with increasing age suggests that it might be unsuitable for this purpose in contexts where the hippocampus undergoes anatomical alterations (Poppenk, 2020). Here, we observed significant age-related variation in the position of the uncal apex, with an estimated annual difference in the anterior direction across the adult lifespan (20-79 years) of approximately 0.038 mm. To circumvent potential variation being introduced by normalization, we also included an additional, native space, approximation of uncal apex location. This measure was the distance in mm from the most anterior point of the hippocampus to the uncal apex, which displayed an annual shortening of approximately 0.02 mm. Alternatively, the relative position of the uncal apex along the hippocampus longitudinal extent could have been used – expressed as the difference in distance between the uncal apex and i) the most anterior point of the hippocampus, and ii) the most posterior point of the hippocampus (Snytte et al., 2022). Whereas Snytte and colleagues (2022) did not observe any age-related variation in this relative measure across the adult lifespan, this estimate should take into account potential age-related decreases in the total length of the hippocampus (in the present study larger than the estimated shortening of the uncus), likely attenuating effects of age on uncal apex position.

Effects of uncal apex position on anterior and posterior hippocampal volume were in line with, although consistently smaller than, those reported previously (Poppenk, 2020). A more anteriorly located uncal apex was linked to smaller anterior and larger posterior hippocampal volumes. Together with an average of 1.4 mm anterior displacement of the uncal apex between younger and older adults, this indicates that not accounting for variation in uncal apex position could lead to the misclassification of approximately 30 mm^3^ (or 1%) of anterior tissue in comparing these groups. The larger estimate of misclassification (∼5%) reported by Poppenk (2020) likely stems from both including a larger sample and, importantly, to the wider age span of participants (covering also the 9^th^ decade).

As expected in the context of uncal apex displacement in the anterior direction, the proportion of anterior hippocampal volume was significantly smaller following landmark-based compared to coordinate-based segmentation. Surprisingly, however, landmark-based segmentation did not lead to overestimation of age-related anterior hippocampal volume loss. Instead, while coordinate-based segmentation showed equal decline in anterior and posterior volumes, landmark-based segmentation showed decline only in posterior volumes. It is possible that the relatively small uncal apex displacement in our sample was insufficient to drive the predicted over– and underestimation of volume loss. Notably, the significant age effect on anterior volume, observed after adjusting for estimated misclassified tissue, suggests that the subuncal area may be a primary contributor to anterior hippocampal volume loss. Additional data from larger samples are, however, needed to comprehensively evaluate a potential critical threshold of uncal apex displacement, as the pattern suggested between subgroups of older adults in our sample was not conclusive (Supporting Figure 1). Furthermore, although gray matter atrophy currently constitutes the main candidate for the underlying source of uncal apex displacement in aging, it is possible that other factors, such as alterations in the shape of the hippocampus (Bussy et al., 2021) may also contribute.

Only landmark-based volumetric findings mirrored the general pattern of greatest posterior hippocampal decline evident across both longitudinal (Chauveau et al., 2021; Langnes et al., 2020) and cross-sectional studies covering the adult lifespan (Hoang et al., 2024; Kalpouzos et al., 2009; Malykhin et al., 2008, 2017; Nordin et al., 2018; Snytte et al., 2022) – despite variation in methods – which include native-space uncal apex landmark-based segmentation (Chauveau et al., 2021; Hoang et al., 2024; Malykhin et al., 2008, 2017; Snytte et al., 2022), segmentation based on the MNI coordinate y –21 (Langnes et al., 2020; Nordin et al., 2018), and voxel-wise analyses (Kalpouzos et al., 2009). Although the opposite pattern (i.e., greatest anterior decline) has been reported (Ta et al., 2012), most of these studies excluded younger and middle-aged adults (Chen et al., 2010; Gordon et al., 2013; Hackert et al., 2002; Nordin et al., 2017), as such capturing the age segment across which longitudinal data seem to suggest a steeper anterior age trajectory (Langnes et al., 2020).

Uncal apex position predicted composite episodic memory performance, as well as word recall performance over and above age and sex, such that a more anteriorly located uncal apex was linked to poorer memory. However, this was not replicated in native-space uncal apex distances. This warrants some degree of caution when evaluating the association between uncal apex position and memory, and necessitates replication in additional samples to establish the source and extent of behaviorally meaningful variation in this normalization-dependent measure.

Consistent with a specialization of function along the hippocampus’ longitudinal axis (Grady, 2019; Persson et al., 2018; Poppenk et al., 2013; Strange et al., 2014), anterior and posterior hippocampal volumes may constitute better predictors of memory than whole hippocampal volume (Hoang et al., 2024; Malykhin et al., 2024; Poppenk & Moscovitch, 2011). Research in young adults emphasize a trade-off between anterior and posterior volume, where higher posterior-to-anterior ratios support episodic recollection (Poppenk & Moscovitch, 2011; Snytte et al., 2020). Adjusting for tissue misclassification due to uncal apex variation, we indeed observed that word recall was predicted by the ratio of posterior to anterior volume. These findings align with longitudinal evidence differentially linking recognition and recollection to anterior and posterior hippocampal volume in aging (Persson & Andersson, 2022), and cross-sectional data linking verbal recollection to the posterior hippocampus (Hoang et al., 2024; Malykhin et al., 2024), but diverge from longitudinal evidence implicating the anterior hippocampus in verbal recollection (Langnes et al., 2020).

While factors beyond differences in segmentation methods may explain inconsistencies in volume-memory associations in aging, our results underscore the importance of maximizing specificity in volumetric estimates, and highlight the importance of accounting for age-related uncal apex differences.

## 5. Conclusion

Using both normalized and native-space estimates, we demonstrate a significant age-related anterior displacement of the uncal apex across the adult lifespan, leading to differences in anterior and posterior volumetric estimates and their associations with age and memory between segmentation methods. Importantly, our findings suggest that normalized y-plane coordinates representing uncal apex position may independently predict memory performance. However, this should be further validated in independent samples and across diverse memory measures. Ultimately, longitudinal data is needed to assess uncal apex displacement during aging and to identify the underlying factors driving this anatomical change.

## Data availability statement

Data from the DyNAMiC study are not publicly available. Access to the original data may be shared upon reasonable request from the Principal investigator, Dr. Alireza Salami.

## Ethical approval

The DyNAMiC study was approved by the Regional Ethical board in Umeå, Sweden (2017/248-31).

## Acknowledgements

This work was supported by grants awarded to A.S. by the Swedish Research Counsel (grant number 2016-01936), Knut and Alice Wallenberg Foundation (Wallenberg Fellow grant), Riksbankens Jubileumsfond (P20-0515), StratNeuro grant at Karolinska Institutet, and to K.N. by Karolinska Institutet Research Foundation (2022-02366). Freesurfer calculations were enabled by resources provided by the Swedish National Infrastructure for Computing (SNIC) at HPC2N (Umeå), partially funded by the Swedish Research Council through grant agreement no. 2018-05973.

