## Supporting Information for "Evaluating the position of the uncal apex as a predictor of episodic memory across the adult lifespan"

### Contents

Effect of segmentation method on anterior hippocampal volume in older age.......................... 2
Supporting Table 1. *Associations with object-location and number-word recall subtests* ........ 3

**Effect of segmentation method on anterior hippocampal volume in older age**

The observation that anterior hippocampal volume from landmark-based segmentation remained stable with age, while coordinate-based anterior volumes significantly declined as a function of age, stands in contrast to the prediction that age-related anterior displacement of the uncal apex would lead to an over-estimation of anterior hippocampal volume loss. To better understand the effect of segmentation method on anterior hippocampal volume in older age, we categorized older individuals (age ≥ 60 years, n = 56) into three subgroups based uncal apex position tertiles. A repeated measures ANOVA tested a potential subgroup-by-segmentation interaction effect on anterior hippocampal volume. The interaction effect was not significant ($F = 1.109$, $p = 0.338$), although the pattern of group differences between segmentation methods suggested that older adults with the most anterior uncal apex position (at, or more anterior to, y = -17.5) only showed numerically lower anterior hippocampal volume compared to the other older subgroups following landmark-based segmentation, Supporting Figure 1.

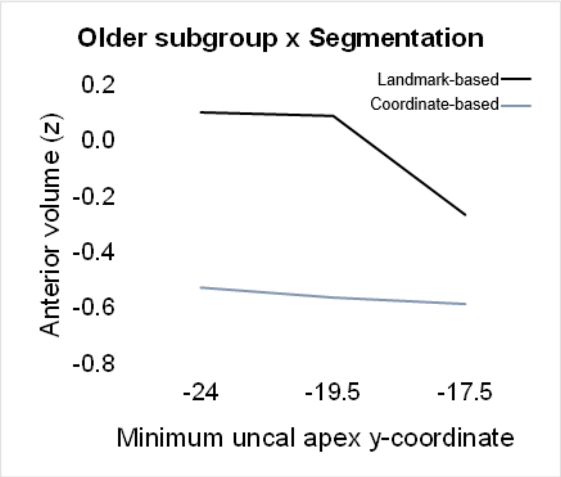


*Supporting Figure 1.* Anterior hippocampal volume from landmark- and coordinate-based segmentation in older subgroups defined based on uncal apex position.

Supporting Table 1. *Associations between hippocampal measures and memory*

| | Episodic composite | Word recall | Object-position recall | Number-word recall |
|---|---|---|---|---|
| Uncal apex | | | | |
| Position | -0.153* | -0.208** | -0.017 | -0.134^ |
| Distance | 0.097 | 0.063 | 0.026 | 0.119 |
| Landmark-based | | | | |
| Anterior volume | 0.066 | -0.033 | 0.034 | 0.135^ |
| Posterior volume | 0.054 | 0.102 | 0.033 | -0.004 |
| Coordinate-based | | | | |
| Anterior volume | 0.012 | -0.071 | 0.010 | 0.080 |
| Posterior volume | 0.033 | 0.037 | -0.006 | 0.051 |
| Posterior:anterior ratio | | | | |
| Landmark-based | 0.004 | 0.095 | 0.002 | -0.099 |
| Coordinate-based | 0.036 | 0.125 | -0.015 | -0.016 |

Partial correlations (Pearson's r) controlling for age and sex; $^{\wedge}p < 0.085$, $^{*}p < 0.05$, $^{**}p < 0.01$
